# Koala retrovirus dynamics in the Snowy Monaro, NSW

**DOI:** 10.64898/2026.10.29.762995

**Authors:** Michaela D. J. Blyton, Karen Marsh, Murraya R. Lane, Arianne Lowe, Keith Chappell

## Abstract

Koala Retrovirus (KoRV) is in transition from exogenous virus to endogenous element and shows distinct profiles between northern and southern koala populations. Northern animals carry high proviral copy numbers of endogenous KoRV-A and a range of exogenous subtypes, while southern koalas instead generally only possess degraded and recombinant versions of the virus. In this study we investigated the KoRV profiles of koalas in the Snowy Monaro (SM) region of southern NSW, which is at the intersection between northern and southern populations. We found that SM koalas have lower KoRV loads and proviral copy numbers than koalas from other northern populations. Plasma RNA subtype profiles were very consistent between individuals with all animals having KoRV-A at low relative abundance and high proportions of subtypes D and L. This pattern of subtype profiles is unique compared to other populations studied and suggests that KoRV-D and/or L may be endogenous in southern NSW or that KoRV-A is effectively suppressed in koalas from the region. Despite these unique profiles, in agreement with findings from other regions, higher KoRV loads were associated with *C. pecorum* shedding. Further, KoRV loads were temporally stable with no consistent changes found over time in koalas in the wild or koalas rescued from burnt habitat after fire and brought in for rehabilitation. Further studies into the role KoRV plays in disease susceptibility in southern NSW and into the endogenization status of KoRV-D/L will provide important insights into this model of retroviral endogenization and guide koala conservation efforts in the region.

**Importance:** Degraded endogenous retroviruses constitute over half of the genetic makeup of all vertebrates, including humans. While most endogenous retroviruses are ancient, koala retrovirus (KoRV) is currently invading the koala genome and still exists as a disease-causing virus. In this study we investigated the KoRV profiles of koalas in southern NSW, which is at the intersection of northern and southern koala populations that show differing KoRV patterns. We reveal that the region has unique KoRV profiles compared with other areas, although KoRV was still temporally stable and associated with susceptibility to Chlamydial infection as seen in other regions. Importantly, we found evidence that additional subtypes (D and L) beyond KoRV-A may be endogenous in southern NSW, providing compelling insights into the ongoing process of KoRV endogenization. This work not only advances our understanding of this model system of retroviral endogenization but also informs conservation efforts for the now endangered koala.

**Data Summary:** Amplicon sequencing data for the KoRV *env* hypervariable region used to identify subtype carriage can be accessed via the NCBI SRA database under BioProject PRJNA1552261. For koala metadata, KoRV load and KoRV subtyping profiles see supplementary material Table S1

## Introduction

Koalas (*Phascolarctos cinereus*) in the north-eastern Australian states of New South Wales (NSW) and Queensland (QLD) are now listed as endangered as a result of population declines due to the combined threats of habitat loss and disease (1). By contrast, koala populations in the southern Australian states of Victoria and South Australia are stable or expanding and, in some regions, koala densities have exceeded carrying capacity, leading to over browsing (2–4). The reason for this dichotomy is likely multifactorial, however, it has been well documented that koala populations in the northern states are heavily impacted by *Chlamydia pecorum* infection that threatens population viability through high rates of infertility and mortality (5). Additionally, we have previously shown that high koala retrovirus load likely increases susceptibility to *Chlamydia* infection and disease in Queensland koalas (6, 7). Notably, koala retrovirus (KoRV) is endemic throughout northern koala populations, existing as an endogenous virus (KoRV-A) integrated into the koala germline, and as a wide diversity of presumed exogenous subtypes (KoRV-A-M) that are primarily acquired from the mother prior to independence (8–10). By contrast, the presumed exogenous KoRV subtypes (B-M) appear to be absent from the southern states and the endogenous form is generally a defective recombinant (recKoRV) and at low copy number (11–14). This suggests that koala retrovirus dynamics may be contributing to the stark differences in koala populations between the northern and southern Australian states (13). As such, investigating KoRV dynamics at the boundary between northern and southern koala populations is pertinent to koala conservation at both the regional and national levels.

The NSW Snowy Monaro koala population is located to the north of the Victorian boarder (Figure 1) and possesses distinct KoRV profiles compared with other NSW populations (11, 12). This contrasts with koala populations on the Victorian side of the border that are genetically homogenous with other koala populations further south, likely due to past population bottlenecks and repeated translocations (11, 12, 15). In a previous genetic analysis of the geographic distribution of the KoRV subtypes, we found that the Snowy Monaro population had (1) higher non-KoRV-A sequence diversity compared to all other populations; (2) comparable KoRV-A sequence richness to southern populations (with both higher than northern populations); (3) divergent KoRV-D sequences; and (3) a unique subtype (KoRV-L) not found in any other population (11). We and others have also found that Snowy Monaro koalas have lower *pol* proviral copy numbers per cell compared to other northern koala populations based on qPCR of the KoRV *pol* and koala beta actin genes from faecal DNA (11, 12). Together these findings suggest that the KoRV profiles of the Snowy Monaro koala population are a mixture of NSW and Victorian profiles but with additional unique features to the region. This is consistent with a recent genomic study of koalas across their range that showed that the Sydney basin forms a biogeographic barrier for koalas, leading koalas in southern NSW (including the Snowy Monaro) to be genetically distinct from those in northern NSW (15). The same study also found that the Snowy Monaro population has some prior admixture with Victorian koalas, although there is no evidence of current geneflow (15). Given the clear uniqueness of the region’s koalas, further investigation of how key KoRV parameters are implicated in koala disease susceptibility is warranted. As koala retrovirus is in transition from an exogenous disease causing virus to degraded endogenous element (16, 17), its association with koala disease is complex and multidimensional. Endogenous KoRV-A expression may be proinflammatory (18) and particular KoRV-A insertion sites are associated with neoplasia (19, 20). In contrast, exogenous KoRV-B carriage was initially associated with chlamydiosis and neoplasia in southern Queensland koalas and captive koalas in the USA (21–23) with KoRV-E and KoRV-F later also associated with neoplasia (24). Further, our assessment of wild koalas admitted for rehabilitation in south-east Queensland found that the proportion of KoRV load (KoRV *pol* RNA concentration in the plasma) attributed to the exogenous subtypes (B-I) was predictive of disease symptoms (6). Additionally, higher exogenous KoRV proportions were positively associated with higher KoRV loads (6). Early small-scale studies of the association between KoRV load and secondary disease produced mixed findings (17, 25–29). However, our large scale study of Queensland koalas showed that high plasma KoRV loads are strongly associated with chlamydial disease pathologies (6). In a recent temporal study of south-east Queensland and mid-north NSW coast koalas, we showed that KoRV load did not increase in wild koalas when they began shedding *C. pecorum* or decrease when they then tested negative. Instead, koalas that were treated for chlamydiosis maintained higher KoRV loads than their healthy counterparts. This suggests that higher KoRV loads increase koalas’ susceptibility to infection and disease (7), potentially through suppression of the adaptive immune system (30, 31). Yet, given the unique KoRV profiles of southern NSW koalas, it is unclear if these same dynamics hold for the Snowy Monaro and the virulence of divergent KoRV-D and KoRV-L is unknown.

**Figure 1:**
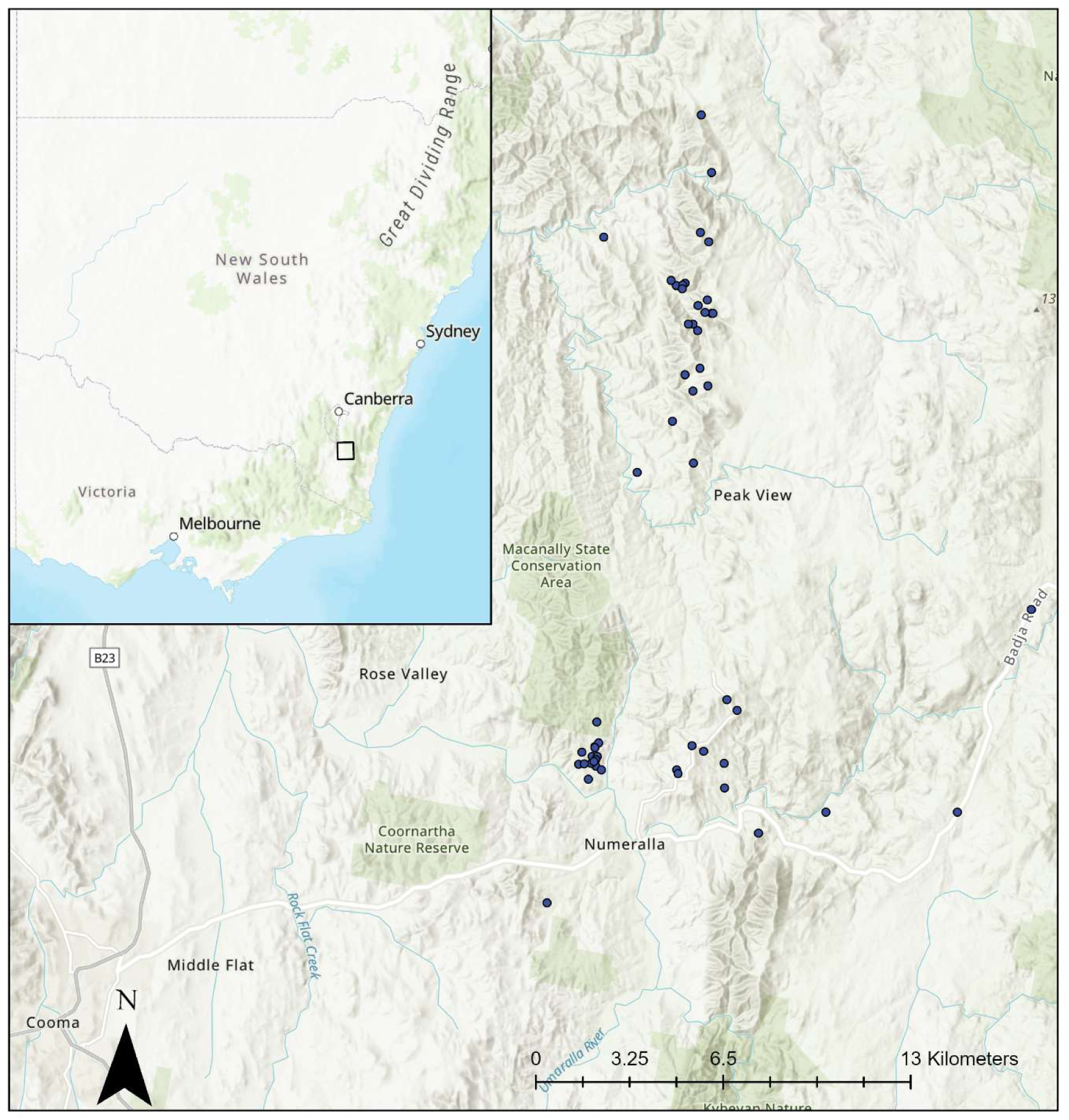
Map showing the geographic coordinates of koalas in this study (blue points) when samples were collected (excluding the koala from Nerriga). For samples that were collected immediately before koalas in rehabilitation were released, the location is given as the tree they were released into. The box in the top right insert indicates the area shown in the main map.

In this study we aimed to investigate the KoRV dynamics of Snowy Monaro koalas, with a particular focus on parameters that are most pertinent to koala disease susceptibility. Specifically; 1) we characterised and assessed the associations between KoRV proviral copy numbers, KoRV plasma RNA load and subtype profiles across 47 koalas from the Snowy Monaro region of the NSW; 2) we assessed the association between KoRV load, koala demographics and *C. pecorum* shedding; and 3) we assessed the stability of KoRV load over time in a subset of 16 koalas.

## Results

### KoRV profiles in NSW Snowy Monaro koalas

The proviral *pol* copy number per cell, as estimated from buffy coat DNA from the 30 rescued koalas, ranged from 7.9 to 23.5 with a mean of 15.9. Log10 *pol* RNA copies per ml of plasma (KoRV load) ranged from 3.23 to 7.36 with a mean of 5.19 among the first samples collected from all 47 koalas. There was no significant association between proviral copy numbers and KoRV load (t = -1.31, p = 0.20). KoRV load showed a bimodal distribution with most koalas having loads of < 5.5 with a mode of between 4 and 4.5, while a smaller number of koalas had loads >5.5 with a mode between 6 and 6.5 (Figure 2).

**Figure 2:**
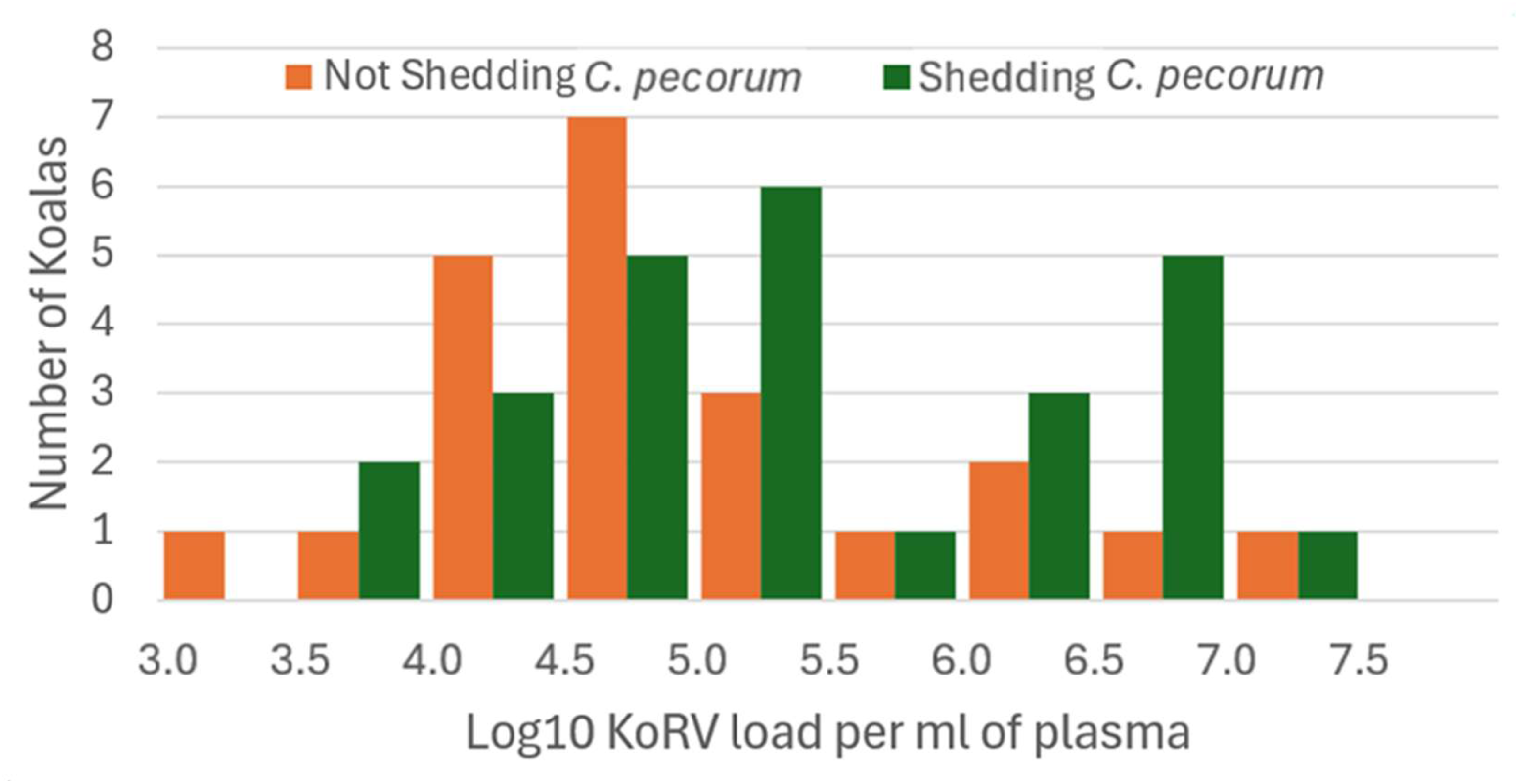
Log10 KoRV load of NSW Snowy Monaro koalas at first sampling, separated by whether the koalas tested positive for *C. pecorum* using qPCR of urogenital and/or ocular swabs.

KoRV-A was detected in the plasma RNA of all 30 rescued koalas at a relative abundance of < 27% in 24 and >97% in five of the koalas (Figure 3). KoRV-D was detected in 28 of the koalas at an average relative abundance of 32.15%. KoRV-L was also detected in 26 of the koalas at an average relative abundance of 38.51%. No other subtypes were detected in any of the sampled koalas, although non-functional sequences were detected in 20 of the koalas at very low relative abundance. These non-functional sequences were at lower relative abundance in the SM koalas than previously reported for QLD koalas, however this can be attributed to a bioinformatics processing difference between this study and our initial study of koalas brought into Currumbin Wildlife Sanctuary for rehabilitation (6).

**Figure 3:**
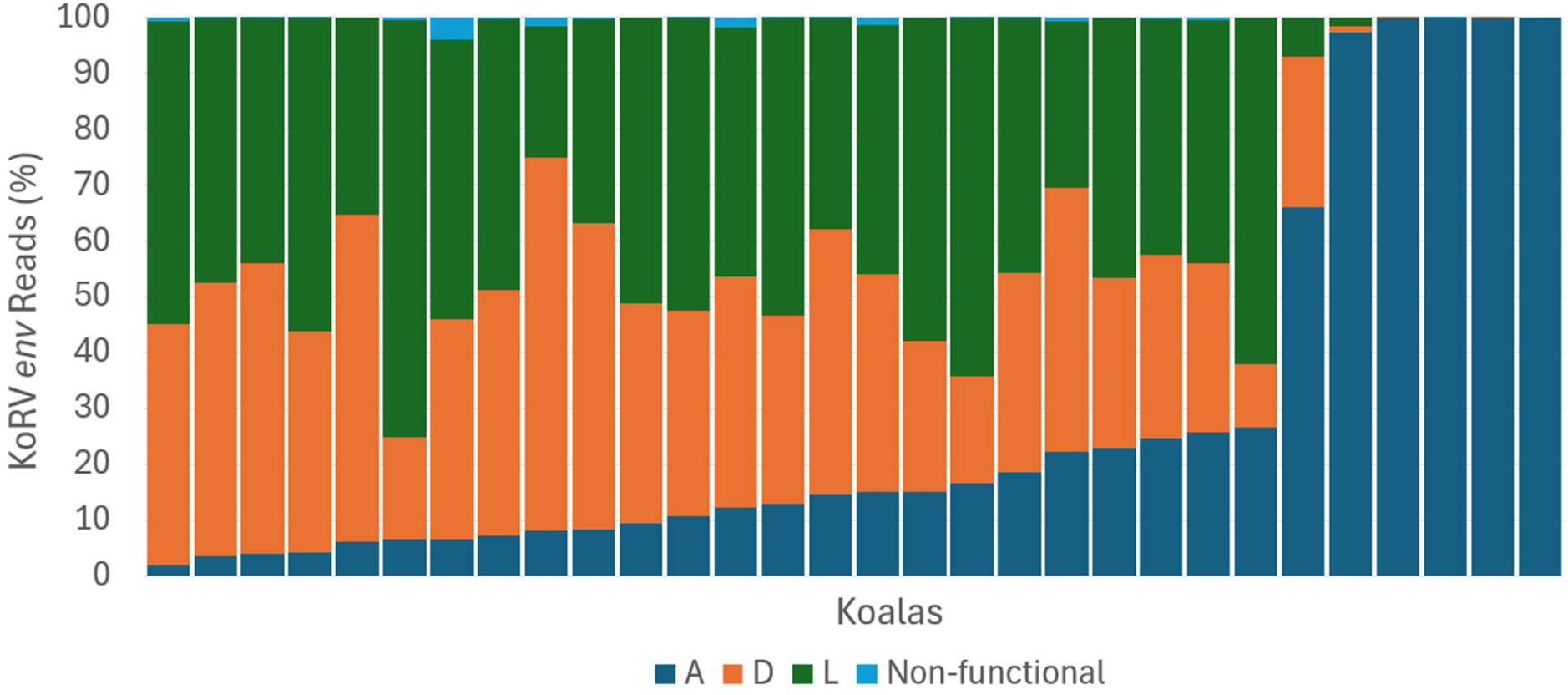
Relative abundance of the KoRV subtypes in plasma RNA of Snowy Monaro koalas as estimated from deep sequencing of the hypervariable region of the KoRV *env* gene.

Notably, all six koalas with a KoRV-A relative abundance of >27% had KoRV loads of 5.52 or less. Although KoRV-A relative abundance and KoRV load were not significantly associated, this was likely due to small sample size and the bimodal distribution of each variable (t = -1.698, p = 0.101).

### KoRV load associations

KoRV load was significantly higher in koalas in the older tooth wear classes (t = 3.22, p = 0.002; Figure 4). KoRV load at first sampling did not differ between *in situ* koalas in burnt or unburnt habitat or *ex situ* rescued koalas (p = 0.746). KoRV load also did not differ between the sexes (females = 24, males = 23; p = 0.428) or with body condition score (p = 0.893).

**Figure 4:**
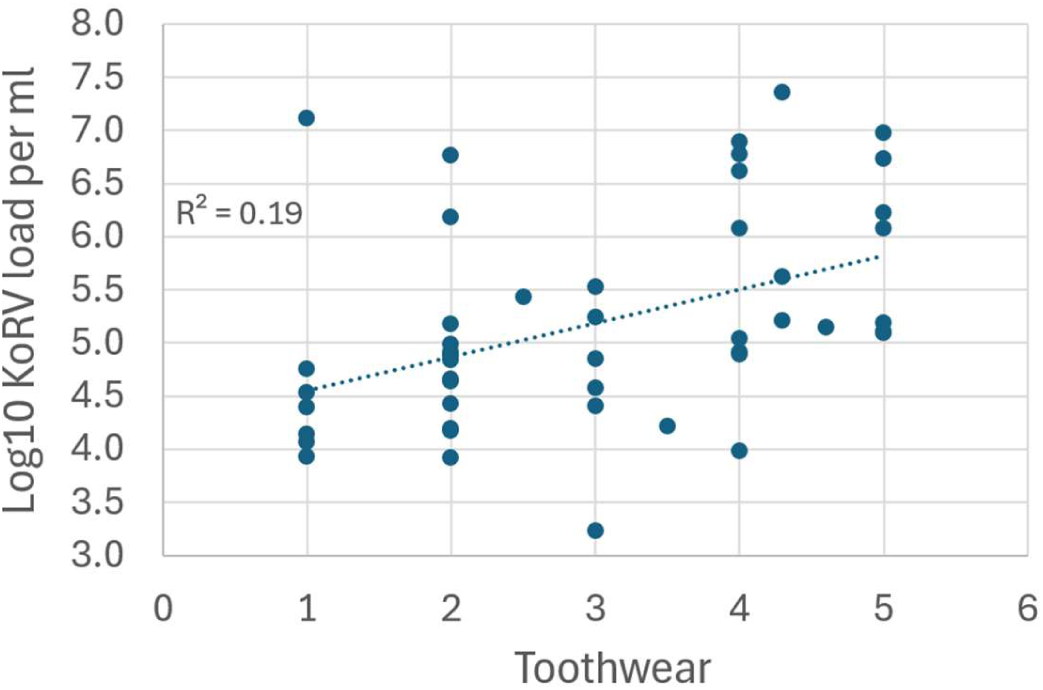
KoRV load in Snowy Monaro koalas at first sampling by tooth wear class, which is correlated with koala age.

Eight of the 47 koalas had rump scores greater than zero on initial assessment. However, all but one of those was scored as one, which corresponds to mild rump discolouration and is often the result of other causes besides *Chlamydia* infection. Additionally, only three koalas tested positive on qPCR for ocular *Chlamydia* infection. Thus, statistical analysis of the association between KoRV load and external signs of disease or ocular *Chlamydia* infection was not conducted. Twenty-six koalas were qPCR positive for urogenital *Chlamydia* infection, with one concurrently positive for ocular *Chlamydia.* Koalas that were positive for *Chlamydia* infection at either body site had significantly higher KoRV loads than koalas negative at both body sites (one-tailed t-test: p = 0.035; Figure 2).

### Temporal patterns of KoRV load

There were no consistent changes in KoRV load among the rescued koalas during their time in captivity with slight increases seen in five and slight decreases in three (Figure 5a). Similarly, there were no consistent changes in KoRV load over time in koalas in the wild, neither for the koalas in burnt or unburnt habitat nor the rescued koalas after release (Figure 5). The average change in KoRV load among the rescued koalas was 0.46 ± standard deviation: 0.41, while for the wild koalas it was 0.31 ± 0.30.

**Figure 5:**
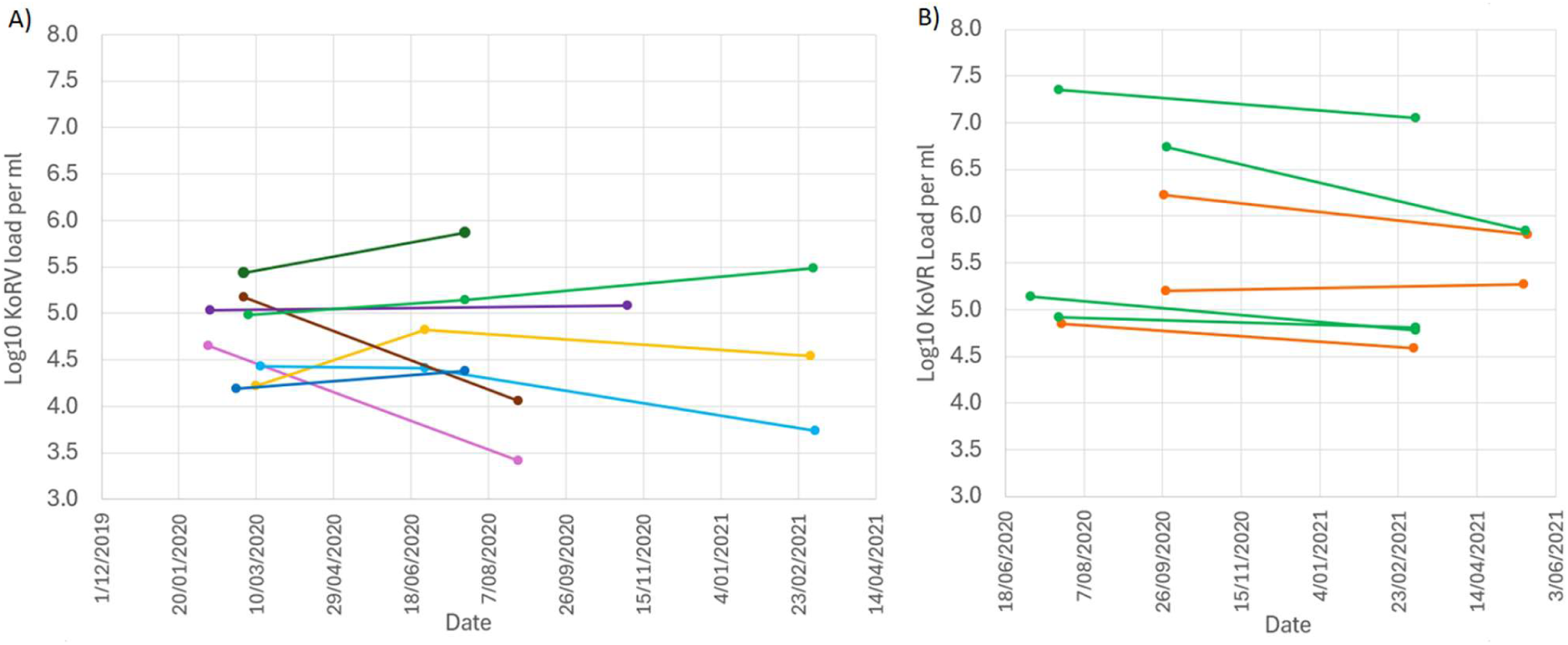
KoRV load in Snowy Monaro koalas over time. Each point represents a sample, with lines joining samples from the same koalas. A) koalas rescued from burnt habitat after the Black Summer fires, with the first sample taken from each koala shortly after rescue, the second sample taken prior to release and the third sample taken in the wild. B) koalas collared and radio tracked in burnt (orange) and unburnt (green) habitat.

## Discussion

In the first assessment of KoRV load and plasma subtype proportions in a southern NSW population, we have revealed that, on average, wild Snowy Monaro (SM) koalas have lower KoRV loads (mean = 5.19) than other wild northern populations studied, including healthy koalas from the Gold Coast and koalas from Moreton Bay in South-East Queensland (GC: mean = 6.10; MB: mean = 7.13) (7, 29). Proviral *pol* copy numbers were also lower in SM koalas than observed in other northern populations, consistent with our analysis of faecal samples from across the koala’s range (32), though interestingly, the two measures were not correlated at the individual level. This suggests the additional role of other factors in driving plasma KoRV RNA load. For instance, we have recently shown that average faecal glucocorticoid metabolite levels are associated with KoRV load, suggesting a role of stress in modulating long term KoRV expression (7). Further, we previously found that KoRV-A proportions and KoRV load are negatively correlated in GC koalas (6), suggesting that the exogenous subtypes increase overall expression levels. Consistent with that finding, the five SM koalas with KoRV-A proportions over 97% in this study had KoRV loads less than 5.5 and were therefore in the low peak of the bimodal distribution of KoRV loads. However, we also found that overall, SM koalas had lower proportions of KoRV-A (mean = 28.9%) compared with either captive mid-north coast NSW koalas (61.3%) or GC koalas (mean = 35.4) despite captive mid-north coast NSW koalas having similar KoRV loads and GC koalas having higher. The low KoRV-A to KoRV-D/L proportion in the absence of high KoRV load in the Snowy Monaro is unusual and different from what has been reported from any other area studied to date.

Looking at this further, we also found that the KoRV-A load was lower than observed in other populations. In fact, SM koalas on average had a quarter of the KoRV-A plasma RNA per *pol* proviral copy compared with GC koalas (mean non-logged KoRV load x mean KoRV-A proportion / mean proviral copy number). In contrast, KoRV-A plasma expression was similar between the mid-north coast and the Snowy Monaro per *pol* proviral copy due to higher proviral copy numbers in northern NSW. Some koala populations have been found to carry a KoRV insertion that triggers the piRNA system in the koala testes, leading to effective suppression of KoRV transcription during spermatogenesis (33). While this system is unlikely to be directly responsible for plasma KoRV RNA load differences between individuals or populations, it highlights that genetic differences between koala populations due to limited dispersal and positive spatial genetic structure even in the absence of inbreeding (34, 35) can influence KoRV activity. As such, further investigation of the genetic determinants of endogenous KoRV-A suppression in NSW koalas may prove valuable.

Most of the koalas in this study had very similar plasma subtype profiles with high proportions of both KoRV-D and KoRV-L and the absence of any other subtypes. The high relative abundance of KoRV-D and KoRV-L is consistent with the faecal proviral subtype profiles found for koalas in the region (11). As discussed above, the high proportions of these subtypes in the plasma may be due to partial suppression of endogenous KoRV-A and low KoRV-A proviral copy numbers. Although, the very consistent profiles between koalas also raises the possibility that KoRV-D and KoRV-L are in fact endogenous in southern NSW. To investigate this possibility, further analysis of endogenous KoRV integrations via long read genome sequencing of koalas from the region will be needed. However, in either case, the high proportion of KoRV-D/L in SM koalas does not appear to be the result of high activity of exogenous KoRV. As such, the proportion of non-KoRV-A subtypes is unlikely to be strongly associated with susceptibility to secondary disease in the Snowy Monaro region of NSW, in contrast to koalas from the Gold Coast, Queensland (6). Some of the other non-KoRV-A subtypes have been associated with increased susceptibility to neoplasia and secondary diseases independently of their impact on KoRV load (6, 21–24). However, the fact that very high KoRV loads (>8) were not observed in this study and that there was a low incidence of external chlamydial pathology despite high rates of *C. pecorum* shedding may indicate that southern NSW KoRV-D and KoRV-L are less of a concern than other non-KoRV-A subtypes. Although, it should be noted that only 8 of the 47 koalas in this study received a urogenital ultrasound, which can detect pathology not apparent on external exam. Nonetheless, the apparent absence of exogenous subtypes previously associated with higher disease susceptibility and in particular KoRV-B and F from the region is reassuring.

Despite the low rate of external chlamydial disease pathologies in this study, we did reveal that koalas that were shedding *C. pecorum* had significantly higher KoRV loads than koalas that tested negative. This is consistent with our previous studies in the Gold Coast, QLD, which found that koalas with higher KoRV loads were more likely to contract *C. pecorum* and that koalas with disease pathologies had higher KoRV loads (6, 7). This suggests that, despite the lower average KoRV loads and copy numbers in the NSW Snowy Monaro, KoRV is nonetheless having an impact on koala health and disease dynamics in the region. Although, it is plausible that the low average KoRV loads in the region may partially explain the lack of progression of *C. pecorum* infection to overt clinical pathologies in this study.

In contrast to our study of koalas undergoing rehabilitation at Currumbin Wildlife Hospital (6), we did not find any correlation between KoRV load and koala body condition score in this study. Given the low rate of disease pathologies in this study, it is likely that body condition was more heavily influenced by environmental factors, such as drought and post fire habitat quality, than koala health when compared to koalas brought in for rehabilitation at Currumbin Wildlife Hospital. This could explain the lack of association with KoRV load and suggests that KoRV load may not have a direct effect on body condition but rather that both are related to disease in the case of koalas at Currumbin Wildlife Hospital. We did however find an association between KoRV load and age, as indicated by tooth-wear. Tarlinton and colleagues (17) previously identified that younger koalas generally have lower KoRV loads. We have also previously found that the number of subtypes carried by a koala increases with age and that koalas with more subtypes have higher KoRV loads (6). Despite this, subtype carriage cannot explain higher KoRV loads in aged individuals in this study due to low subtype diversity. Instead, it may be that older individuals have a decreased capacity to suppress viral activity as occurs in with HERVK in humans (36).

In agreement with Quigley and colleagues (29) and our assessment of KoRV load over time in wild Gold Coast and captive mid-north coast NSW koalas (7), we found little change in KoRV load over time either in the SM koalas rescued from burnt habitat or koalas radio tracked in the wild. The general temporal stability of KoRV load may also explain why we did not see any differences in KoRV load between burnt and unburnt habitat or in response to captivity after rescue. Despite the association between KoRV load and average faecal glucocorticoid metabolite (FGM) levels, there is not a temporal association between changes in FGM levels and changes in KoRV load within koalas (7). Thus, while the black summer fires, rescue and reduced food resources may be reasonably expected to have produced short-term physiological stress to the koalas, this was not reflected in short-term KoRV load changes.

The Snowy Mountains region of southern NSW is at the boundary between northern and southern koala populations that show distinct population genetics and koala retrovirus profiles. This study has revealed that the koalas in this region possess unique KoRV profiles that differ from those seen either in southern Australia or in other northern populations. We found that the plasma RNA KoRV subtype profiles were very consistent between individuals in the Snowy Monaro with all animals having KoRV-A at low relative abundance and high proportions of subtypes D and L. Yet SM koalas had lower KoRV loads than other wild northern populations. This pattern indicates that KoRV-D and/or L may be endogenous in southern NSW or that KoRV-A is effectively suppressed in koalas from the region. These findings suggest that the Snow Monaro and potentially other southern NSW populations should be managed as a separate unit to more northern NSW koala populations. Nonetheless, we also found that higher KoRV loads were associated with *C. pecorum* shedding, suggesting that KoRV plays a role in disease susceptibility in southern NSW, similar to other northern populations.

## Methods

### Study Design

A total of 47 koalas were sampled for this study from the NSW Snowy Monaro ( Figure 1; Table S1; with the exception of one koala, Belle, who was rescued from Nerriga) that were a subset of those included in a study of post fire health and movement patterns in the region (37, 38). Thirty of the koalas were rescued from burnt habitat during or within one month after the 2019/2020 black summer fires and kept in care for 3 to 9 months until they had recovered and sufficient vegetation regeneration had occurred in their home ranges for them to be released. Most rescued koalas were free from injury, with only one presenting with minor burns. The koalas were initially sampled soon after rescue with eight koalas also sampled prior to release and a third sample taken from two koalas 9 months post release to assess temporal patterns in KoRV load.

The remaining 17 koalas were captured in burnt (n = 6) or unburnt (n = 11) habitat, fitted with radio tracking collars, and released on the same day to investigate their home range size and movement patterns (37). Samples were taken from all koalas when the collars were fitted. A sample was also taken when the collars were removed from seven of the koalas (burnt = 3; unburnt = 4) to assess temporal trends in KoRV load.

### Health assessments

Health checks were conducted by wildlife veterinarians under heavy sedation or general anaesthesia as described in Lane et al. (38). Koalas were checked for the presence of burns or any other clinically important findings, and a variety of general parameters were recorded, including sex, age (from tooth wear score) and weight. Tooth wear score was determined by assessing the wear of the fourth upper premolars as per (39). Body condition was assessed using a standardised technique of palpation of the muscles on either side of the protruding ridge of bone on the scapula (spine of the scapula), and was scored out of 5 (26).

### Sample collection

Blood was collected from the cephalic vein into EDTA tubes for KoRV load, proviral copy number and subtyping analysis. After collection, blood was transported to the Australian National University, centrifuged at 6500× g for 10 min and the plasma and buffy coat transferred into separate tubes and frozen at either -20 or -80 °C within 8 hrs of collection. The plasma from 14 samples were separated into two tubes, one of which was kept at -20 °C for 2 years while the other was stored at -80 °C. Assessment of KoRV load for these duplicate samples by qPCR (see below) showed no significant differences between the storage conditions (paired t-test: p = 0.106) with KoRV load higher in eight of the -20 °C replicates and lower in the remaining six. As such, it was concluded that KoRV load is stable in plasma kept at either temperature and that the samples from the different storage conditions could be reliably compared.

Ocular swabs were collected by gently and carefully rolling a swab (premoistened with sterile saline) across conjunctiva on both eyes, while urogenital swabs were collected from the urogenital sinus (females) or penile urethra (males). Swabs were stored at ambient temperature for up to five days, pending analysis at the School of Veterinary Science, University of Sydney. Chlamydial detection was conducted using an established qPCR (40) for chlamydial 23S, ompB and koala beta-actin as a sample quality control, following DNA extraction on a magnetic bead-based 96-well extraction system (Kingfisher Flex, Thermo Fisher Scientific) with positive and negative controls.

### KoRV proviral copy numbers

Total genomic DNA was extracted from a single buffy coat sample from each of the thirty rescued koalas using the FavorPrep blood genomic DNA mini kit (Favorgen Biotech Corp), as per the manufacturer’s instructions. Buffy coat samples were not available for the 17 radio-tracked koalas. The number of KoRV gene copies per koala cell was estimated for each sample from the number of KoRV *pol* and β-actin molecules per qPCR reaction as per Blyton et al. (32).

### KoRV plasma RNA loads

Total RNA was extracted from all plasma samples using the High Pure Viral Nucleic Acid Kit (roche) according to the manufacturer’s instructions with modifications as outlined in Blyton et. al. (32). First strand cDNA was created using Superscript III (Invitrogen) as per Blyton et. al. (32). KoRV plasma RNA loads were estimated by quantitative PCR of a 110 bp fragment of the KoRV *pol* gene (41) amplified in triplicate from the plasma cDNA as per Blyton et. al. (42).

### KoRV subtype profiles

An approximately 500 bp region of the KoRV *env* gene containing the previously identified hypervariable domain (43) was amplified from the total plasma RNA of the rescued koalas and sequenced. The KoRV subtype profiles of the 17 radio-tracked koalas were not assessed as they were only included in the study to assess temporal variation in KoRV load. The *env* hypervariable region was first amplified using the nested polymerase chain reaction (PCR) protocol described in Blyton et al. (32). Sequencing libraries were prepared from the *env* PCR products using the Nextera XT sample preparation kit (Illumina) following the manufacturer’s protocol. Each library was uniquely tagged using the Nextera XT indexing primers (Illumina). Paired-end sequencing was performed at the Australian Centre of Ecogenomics, on the Illumina Miseq using the version 3 reagent kit for 300 cycles.

Processing of the raw sequencing reads was completed in CLC genomics workbench 20 as per Blyton et al. (32) except that the minimum read length for retention was set to 400 instead of 450 bp. Rarefied reads were then *de novo* clustered across all samples at 100% similarity (ANI) in QIIME 2 (44). Read clusters with less than 3 reads across the dataset were then discarded. The representative sequences for the remaining 100% clusters were then *de novo* clustered across all samples at 97% similarity in QIIME 2 (44). For each 97% ANI read cluster, the most abundant sequence was identified and designated the representative sequence for that cluster. The representative sequences for the 97% ANI clusters were then blasted using default parameters against the NCBI ‘nt’ database to confirm their identity as koala retrovirus *env* sequences.

The representative sequences for the 97% ANI clusters were then translated *in silico* using SeqKit (45). Sequences containing missense mutations, frameshift mutations or large deletions (>4 amino acids) were then identified and designated as non-functional. The translated sequences were then aligned with reference sequences for each of the previously identified subtypes (KoRV-A (46); KoRV-C (AB828005;47); KoRV-E (48); KoRV-B21, D35, F9, G4, H2, I4 (43); KoRV-K (9); KoRV-L and KoRV-M (11)) with MAFFT (49) and visually examined to determine their subtype designation. The presence and abundance of each of the subtypes in each of the samples were then inferred from the representative sequence by sample table generated from the 100% QIIME 2 clustering.

### Statistical Analysis

Associations between KoRV load and other variables were assessed by linear fixed effects models using the lm function in R (50). KoRV load was fitted as the response variable in the analysis with either proviral copy number, plasma RNA KoRV-A proportion, tooth wear, sex, body condition score or cohort (burnt, unburnt or rescued) fitted as the explanatory variable. Only the first sample from each koala was included in the analysis. Due to the relatively small sample size of the study, multivariate models were not considered. The assumption that the model residuals conformed to normality was assessed by Shapiro-Wilks test. The residuals of the model where body condition score was the explanatory variable did not conform to normality unless KoRV load was first rank transformed. All other models conformed to normality.

To assess if KoRV load differed between koalas that tested positive or negative for *C. pecorum* infection a one tailed, two sample t-test with homeostatic variance was conducted. A one-tailed test was chosen as previous studies have indicated that koalas with *C. pecorum* infection have higher KoRV loads (6, 7).

## Ethical Approval

Sample collection was approved by the NSW Government Department of Planning, Industry and Environment Animal Ethics Committee (permit number 200421/04) and was conducted in accordance with the Australian Code for the care and use of animals for scientific purposes.

## Funding information

Sample collection was funded by the NSW Government under the NSW Koala Strategy and the Australian Government Wildlife and Habitat Bushfire Recovery Program (DOC20/500625); and through donations from Two Thumbs Wildlife Trust and the Minderoo Foundation (TI_ANU001). The molecular analysis was funded by the Australian Research Council (DP180103362).

## Acknowledgements

We thank Mariska de Buin for completing the KoRV load qPCRs and Julian Zaugg for completing the KoRV subtyping bioinformatic analysis. The *C. pecorum* qPCR testing was conducted by the Koala Health Hub at the University of Sydney. We also thank Dr Jelena Vukcevic and Dr Larry Vogelnest for conducting some of the veterinary work associated with sample collection.

## Author Contributions

The project was conceptualised by MB and KC. ML, AL and KM collected the samples, and AL undertook most veterinary assessments of the koalas. MB undertook the KoRV laboratory work. MB analysed the data and wrote the manuscript. KC provided the funding for the KoRV analysis. All authors edited the manuscript and approved the final version. We acknowledge the peoples of the Ngarigo Nation, who are the traditional owners of the lands on which this study was conducted.

## Conflicts of Interest

The authors declare that there are no conflicts of interest.

